# Highly accurate genome assembly of an improved high-yielding silkworm strain, Nichi01

**DOI:** 10.1101/2022.11.14.516399

**Authors:** Ryusei Waizumi, Takuya Tsubota, Akiya Jouraku, Seigo Kuwazaki, Kakeru Yokoi, Tetsuya Iizuka, Kimiko Yamamoto, Hideki Sezutsu

## Abstract

The silkworm (*Bombyx mori*) is an important lepidopteran model insect, and an industrial domestic animal traditionally used for silk production. Here, we report the genome assembly of an improved Japanese strain Nichi01, in which the cocoon yield is comparable to that of commercial silkworm strains. The integration of PacBio Sequel II long-read and ddRAD-seq-based high-density genetic linkage map achieved the highest quality genome assembly of silkworms to date; 22 of the 28 pseudomolecules contained telomeric repeats at both ends, and only four gaps were present in the assembly. A total of 452 Mbp of the assembly with an N50 of 16.614 Mbp covered 99.3% of the complete orthologs of the Arthropod core genes. Although the genome sequence of Nichi01 and that of the previously reported low-yielding tropical strain p50T assured their accuracy in most regions, we corrected several regions, misassembled in p50T, in our assembly. A total of 18,397 proteins were predicted using over 95 Gb of mRNA-seq derived from 10 different organs, covering 96.9% of the complete orthologs of the Arthropod core genes. The final assembly and annotation files are available in KAIKObase (https://kaikobase.dna.affrc.go.jp/index.html) along with a genome browser and BLAST searching service, which would facilitate further studies and the breeding of silkworms and other insects.

## Introduction

The domestic silkworm, *Bombyx mori* (Lepidoptera: Bombycidae), has been used as a primary resource in silk production systems for thousands of years. Its superior ability to secrete protein is utilized not only for fiber production but also for the production of recombinant proteins for medical applications, which increases the need to understand the genetic mechanisms of efficient silk production in silkworms (Sezutsu *et al.* 2018). In 2004, the first reported genome assembly of the silkworm strain p50T provided the basic information required for its genetic analysis (Mita *et al.* 2004; Xia *et al.* 2004). The genome sequence of p50T was improved in 2008 by integrating two different assemblies of the strain that were reported independently by Chinese and Japanese research groups (The International Silkworm Genome Consortium, 2008). In 2019, genome sequences of p50T were assembled using PacBio long reads and BAC sequences, which delivered a highly contiguous assembly (Kawamoto *et al.* 2019). These assemblies contribute to genetic research, thereby advancing our understanding of the molecular mechanisms of silk production, such as the identification of the transcription factor that regulates major silk proteins and the genes associated with cocoon yield (Li *et al.* 2017; Li *et al.* 2020; Takiya *et al.* 2016; Tsubota *et al.* 2016).

Recently, the cost of next-generation sequencing (NGS) has decreased dramatically, accelerating the collection of comprehensive genetic information and enhancing the importance of reference genome assemblies. Downstream analysis using NGS is generally based on mapping reads to the reference genome. Since the selection of the reference genome considerably influences results of the downstream analysis, it is necessary to select the most suitable, best quality reference genome for analytical purposes (Valiente-Mullor *et al.* 2021). There is a wide variety of silkworm strains worldwide (Xiang *et al.* 2018). Improvements in the genome assembly of silkworms and the availability of various suitable strains are critical conditions for the development of silkworm research.

Here, we report a new genome assembly of the improved silkworm strain, Nichi01 (alias: J01). This strain shows high silk productivity, in contrast to the tropical low-yielding p50T strain. This genome assembly can be the best genetic reference for commercial silkworm strains and can accelerate our understanding of the molecular mechanisms that enable their superior protein productivity. It also provides the best accuracy to date among all published silkworm genomes, and was constructed by integrating PacBio long-read sequencing and a traditional linkage map that was constructed using double-digested Restriction Associated DNA sequencing (ddRAD-seq) (Peterson *et al.* 2012). All data are available at KAIKObase (https://kaikobase.dna.affrc.go.jp/index.html), with several useful tools for analysis (Yang *et al.* 2021).

## Materials and Methods

### Origin and derivation of Nichi01

According to a previous report, the Nichi01 strain was developed by crossbreeding two improved Japanese strains, KN9 and HN53, with a high shell/whole cocoon ratio (Azuma, 1985). KN9 was derived from the high-yielding strain 010 (Azuma 1985), which was developed by selective breeding that focusd on cocoon shell weight through 55 generations from the European strain Giallo Asocoli (Maruyama 1984).

### Cocoon-associated phenotyping

Silkworm strains, Nichi01 and p50T, were reared on fresh mulberry leaves under a controlled environment of 12 h light/dark photoperiod at 25 °C. Measurements of the cocoon shell weight and shell/whole cocoon weight ratio were conducted a week after cocooning. The shell/whole cocoon weight ratio was calculated by dividing the weight of the cocoon by the sum total of the weight of the pupa and cocoon.

### SNP calling and principal component analysis

The SRA accession numbers of the published sequencing data used for the analysis are listed in Supplementary Table S1. Raw reads were trimmed using fastp v0.20.0 with the following parameters: -q 20 -n 5 -l 100 (Chen *et al.* 2018). The trimmed reads of each strain were mapped using minimap2 v2.17 with default parameters (Li, 2018). The BAM format of the mapping results was filtered to remove duplicated reads using Picard v2.23.4 (http://broadinstitute.github.io/picard/). According to the GATK best practice introduced on the official website (https://gatk.broadinstitute.org/hc/en-us), quality control of mapping results and SNP and INDEL calling were performed using GATK v4.2.0.0 (McKenna *et al.* 2010). The parameters for filtering the VCF data of SNPs were as follows: QD < 2.0, QUAL < 30.0, SOR > 4.0, FS > 60.0, MQ < 40.0, MQRankSum < -12.5, ReadPosRankSum < -8.0. The parameters for filtering the VCF data of INDELs were as follows: QD < 2.0, QUAL < 30.0, FS > 200.0, SOR > 10.0, and ReadPosRankSum < -20.0. Tassel5 was used for principal component analysis using the homozygous SNPs on the coding sequences (Bradbury *et al.* 2007).

### Sample preparations and sequencings

The genomic or transcriptomic sequencing samples were reared on an artificial diet (NOSAN, Yokohama, Japan) under a 12 h light/dark photoperiod at 25 °C. For whole-genomic sequencing, the genomic DNA of the Nichi01 strain was isolated from the silk glands of two male fifth-instar larvae using Genomic-tips 100/g and Genomic DNA Buffer Set (QIAGEN). Long-read sequencing for genome assembly was performed using continuous long-read (CLR) mode of the PacBio Sequel II System (Pacific Biosciences, Menlo Park, California, U.S.A.). Short-read sequencing for assembly polishing was performed using the NovaSeq 6000 Sequencing System; this paired-end sequencing generated reads with a length of 151 bp (Illumina Inc., San Diego, U.S.A.). The RNA of the Nichi01 strain was extracted from 10 tissues of the third-day fifth instar larva: anterior silk gland (ASG), anterior part of the middle silk gland (MSG_A), the middle part of the middle silk gland (MSG_M), posterior part of the middle silk gland (MSG_P), posterior silk gland (PSG), fat body (FB), midgut (MG), Malpighian tubule (MT), testis (TT), and ovary (OV). The partitioning of the silk gland is illustrated in the supplementary information (Supplementary Figure S1). The RNA sequencing was performed by Illumina NovaSeq 6000 Sequencing System in paired-end sequencing generating reads with a length of 151bp. All the library constructions and sequencing were conducted via the outsourcing service of mRNA-seq (Macrogen Japan Corp., Kyoto, Japan). An F_2_ hybrid population of p50 (the original strain of p50T; p50T is a descendant of p50 maintained at the University of Tokyo, which had undergone repeated passage using a single pair to reduce internal genetic diversity) and Nichi01 was used to construct ddRAD-seq libraries. All sampled F_2_ progenies were female. Genomic DNA was extracted from the pupae of each sample using a DNeasy Blood and Tissue Kit (QIAGEN, Hilden, Germany). Two ddRAD libraries were prepared; the restriction enzyme pairs, EcoRI-HF/MspI and PstI-HF/MspI (New England Biolabs, Ipswich Commonwealth, Massachusetts, U.S.A), were used. The library construction was conducted according to a previous study (Uchibori-Asano *et al.* 2019). The libraries were merged into one sequencing library, which was sequenced using Illumina HiSeq 2000 to generate paired-end reads with a length of 101 bp (Macrogen Japan Corp.). The re-distribution of these reads was performed based on the barcode sequences using the script “process_radtags” in Stacks v1.48 (Catchen *et al.* 2011). Information on all sequencing data used here was summarized in Supplementary Table S2.

### Genome assembly and scaffolding by linkage-map construction

Before assembly, the raw reads of the genomic sequences were filtered. The PacBio long reads with an e-value of 0.0 in BLASTn analysis against the reference mitochondrial sequences of silkworm (NCBI Reference Sequence: NC_002355.1) were regarded as mitochondria-derived sequences, and were removed (Altschul *et al.* 1990). Illumina short reads were trimmed using fastp v0.20.0 with the following parameters: -q 20 -n 5 -l 100 (Chen *et al.* 2018). The remaining long reads were corrected and assembled using Canu v2.1.1 with the default parameters except for the one, “genomeSize”, fixing an estimated genome (Koren *et al.* 2017). Input genome size (450 Mb) was determined based on previous studies (Kawamoto *et al.* 2019; Mita *et al.* 2004; Xia *et al.* 2004). Using the Illumina short reads, the obtained contigs were once polished using Pilon v1.23 with the following parameters: --diploid (Walker *et al.* 2014). To assess haplotypic duplication in the assembly and removal of artifactual contigs, Purge Haplotigs v1.1.1 was performed with a mapping result of a quarter of the PacBio raw reads against the contigs (Roach *et al.* 2018). The coverage histogram as a partial output of the pipeline is shown in the supplementary information (Supplementary Figure S2). To remove only artifactual contigs with extremely low coverage of the reads, each parameter that fixes the read depth low and high cutoff, “-l” and “-h,” was set to 5 and 250, respectively. Seqkit v2.2.0 was used for the subsampling of reads (Shen *et al.* 2016).

The ddRAD-seq reads were mapped to the polished contigs using BWA v0.7.17 with default parameters (Li and Durbin 2009). The script “ref_map.pl” in STACKS v1.48 was run with the parameters: -m 3 -S -b 1 -A F2, to identify SNP and to obtain genotype data of 102 F_2_ intercross progenies, two parents, and two F_1_ hybrid parents (Catchen *et al.* 2011). Individuals and markers with more than 20% of missing data were excluded. The markers on the Z chromosome were extracted using the following criteria: (a) the genotype of the female F_1_ parent should be paternal homozygous type, (b) the genotype of the male F1 parent should be heterozygous, and (c) the genotype of the female F_2_ should be maternal or paternal homozygous. The markers on the autosomal chromosomes were also filtered by confirming that the genotypes of the F1 parents were heterozygous. Onemap v2.8.2 was used to construct linkage maps with an LOD score of 3 (Margarido *et al.* 2007). Markers on autosomes and those on Z chromosomes were analyzed separately. Autosomal markers were analyzed using the model for F_2_ populations. Paternal homozygous genotypes of Z-chromosomal markers were converted to heterozygous genotypes for adaptation to the model for backcross populations in Onemap. All markers were confirmed to follow the expected segregation using the chi-square test-based internal function “test_segregation.” To order the markers according to their genetic distances, the function “order_seq” was used with the parameters; subset.search = “twopt,” twopt.alg = “rec,” THRES = 3. Chimeric contigs were detected by the inconsistency of the genetic and physical positions of the markers; some of the markers, even included in the different genetic linkage groups, were located in the same contig. The misassembled points of the chimeric contigs were identified as points with extremely low coverage of the mapped short reads (Supplementary Figure S3). The validity of the correction of the assembly was verified by alignment with the genomic sequence of p50T. The contigs were aligned into pseudomolecules according to the genetic positions of the markers. When adjacent terminal sequences of aligned contigs overlapped, we considered the fragmentation due to excessive parameter stringency in the assembly and connected them. Some of the contigs without genetic markers were inserted into pseudomolecules using the following criteria: (a) the inserted position should be consistent with the alignment result with genomic p50T sequences provided by RaGOO v1.1 (Alonge *et al.* 2019), (b) the contig should have more than 3.5 kbp of overlapping sequence in its terminal region with the inserted region of pseudomolecule, and (c) the mismatch number should be less than 1% of the overlapping region. After the series of scaffolding processes described above, the constructed pseudomolecules were further polished with short reads using Pilon v1.23. The length of each gap region, consisting of consecutive N, was set to 100 bp. To visualize the genetic and physical positions of the markers, a linkage map was re-constructed with the final genome assembly using the same method.

### Prediction and annotation of repeat sequence and coding gene

RepeatMasker v4.1.2 was used to detect repetitive sequences with the transposable element library provided in the Silkworm Genome Research Program (http://sgp.dna.affrc.go.jp/pubdata/genomicsequences.html) (https://www.repeatmasker.org/) in the RMBlast-dependent mode. The resultant repetitive sequences were soft-masked for coding gene prediction. BRAKER2 and StringTie2 were used as primary tools for gene prediction. We ran BRAKER2 with protein and RNA-seq data, using the following parameters: --prg gth --softmasking -- trainFromGth (Brůna *et al.* 2021). The curated 2,323 protein sequences of *Bombyx mori* with IDs starting with “NP_” provided by NCBI RefSeq database, and BAM format of mapping results of 30 mRNA-seq reads from 10 different organs of Nichi01 were used as hints in the prediction. GenomeThreader v1.7.3 was chosen as an alignment tool for the BRAKER2 pipeline to provide the alignment result of the protein sequences with genomic sequences (Gremme *et al.* 2005). HISAT2 v2.1.0 was used to map each RNA-seq read filtered by fastp v0.20.0 (Chen *et al.* 2018; Kim *et al.* 2019). The mapping results of RNA-seq were also used to predict transcript sequences using StringTie2 v2.2.0 (Kovaka *et al.* 2019). The resultant sequences were translated using TransDecoder v5.5.0 (https://github.com/TransDecoder/TransDecoder). The predicted genes generated by BRAKER2 and Stringtie2 with a length greater than 49 amino acids were integrated using EVidenceModeler v1.1.1 (Haas *et al.* 2008).

### Other tools used in the analysis

Evaluation of genome and proteins by Benchmarking Universal Single-Copy Orthologs (BUSCO) v5 with Arthropoda ortholog set: gVolante (Nishimura *et al.* 2017), drawing chromosome illustration: chromomap v0.4.1 (Anand and Rodriguez Lopez, 2022), dot-plot analysis: D-GENIES (Cabanettes and Klopp, 2018), gene annotation: eggNOG-mapper (Cantalapiedra *et al.* 2021).

### Statistical analysis

The Wilcoxon rank sum test was performed using the initially installed stat command “wilcox.test” in R v4.0.3.

## Results and Discussion

### Cocoon traits and genetic features of Nichi01

Nichi01 is an improved Japanese strain with high silk production. Compared to p50T, Nichi01 was distinctly superior in cocoon-associated traits (Figure 1A-C). The cocoon shell weight and shell/whole cocoon ratio recorded in the Nichi01 were 2.8 and 1.8 times that of p50T, respectively. In analyses using a reference genome, information on the genetic background of the material can be useful for selecting the most appropriate one. As previously described, Nichi01 is derived from both Japanese and European strains (Azuma 1985; Maruyama 1984). However, the detailed genetic background of Nichi01 is unknown. To characterize the genotype of Nichi01, we performed SNP-based principal component analysis using the Nichi01 whole-genomic sequencing data and published data for other strains (Figure 1D, Supplementary Table S1). In total, 1,028,933 homozygous SNPs were used for the analysis. Nichi01 was included in the cluster of Japanese improved strains and differed distinctly from p50T (Kawamoto *et al.* 2019), included in the cluster of Indian local strains and wild silkmoths.

**Figure 1:**
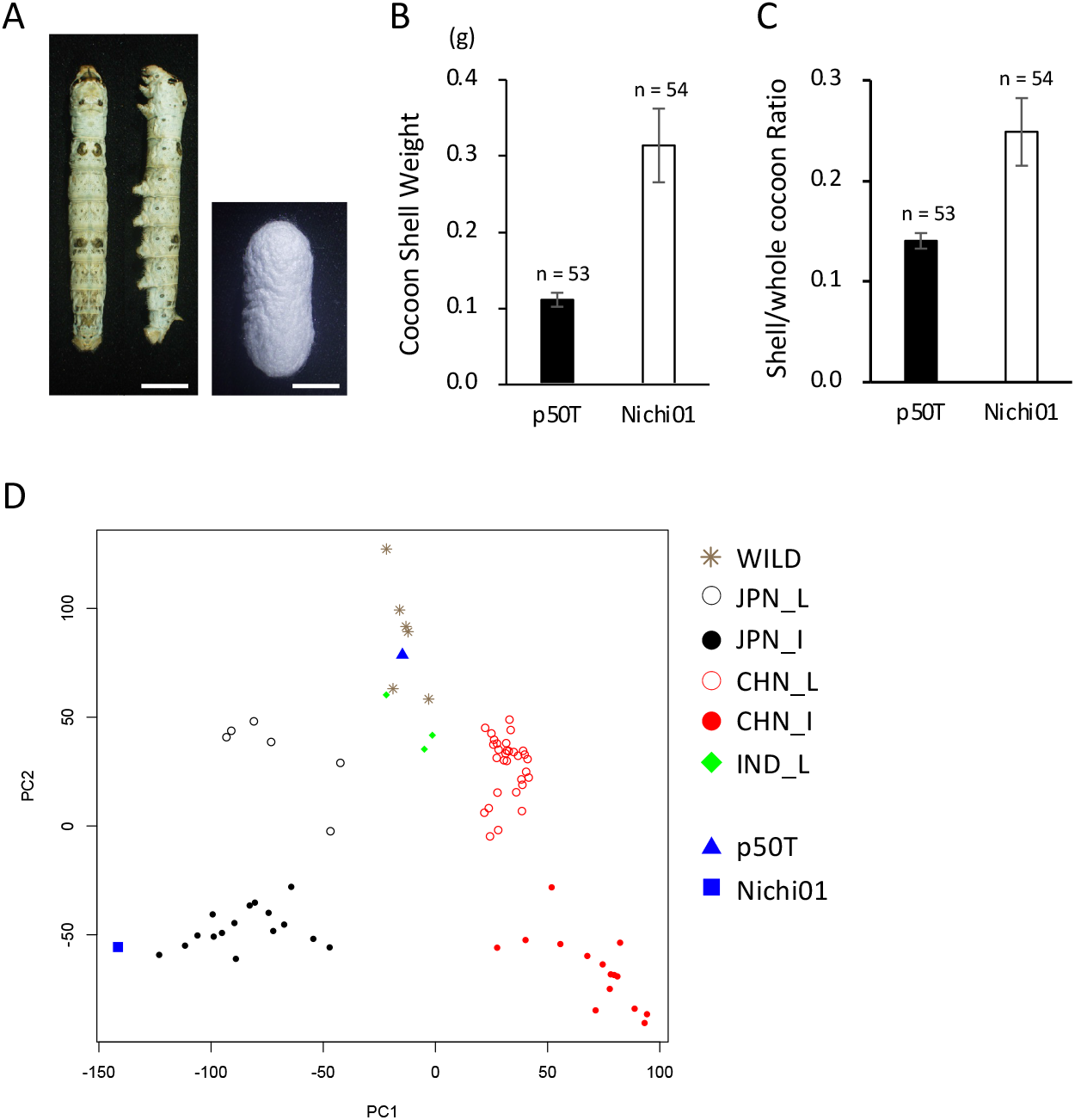
Cocoon-associated traits and genetic property of Nichi01. (A) Larva on third-day of final (fifth) instar (left) and cocoon of male Nichi01 (right). Bar = 10 mm. (B) Cocoon shell weights of male p50T and Nichi01. Data are means ± SD. (C) Shell/whole cocoon ratio of male p50T and Nichi01. Data are means ± SD. (D) Principal component analysis of the genotypes of silkworm strains in the world. The calculation was performed using homozygous SNPs on coding sequences. The sequencing data used here was summarized in Supplementary Table S1 and S2. The labels are according to the previous report (Xiang *et al.* 2018). JPN_I: Japanese improved strain; JPN_L: Japanese local strain; CHN_I: Chinese improved strain; CHN_L: Chinese local strain; IND_L: southern Indian strain; WILD: wild silkmoth *Bombyx mandarina*.

### Genome assembly of Nichi01

To perform *de novo* assembly of the Nichi01 genome, we integrated the three sequencing methods (Figure 2). Sequencing performed by the PacBio Sequel II System generated 166.32 Gb (~373× coverage) of CLR reads with N50 = 14,553 bp for assembly. Sequencing performed using the Illumina NovaSeq 6000 Sequencing System generated 58 Gb (~128× coverage) of paired-end reads with a length of 151bp for polishing. Using Canu (Koren et al. 2017), PacBio long reads were corrected and assembled into 378 raw contigs with a total sequence length of 457 Mb. The raw contigs were polished once using Illumina short reads. Haplotypic duplication in the assembly was assessed using Purge Haplotigs (Roach *et al.* 2018). As Nichi01 is an inbred strain undergoing decades of inbreeding, its genome was presumed to be highly homogeneous; the separative breeding of Nichi01 initiated in 1971 (Azuma 1985). Consistent with the background, the coverage histogram exhibited a single primary peak, indicating that the haploid assembly was almost complete (Supplementary Figure S2). The tools for the removal of duplicated allelic contigs in assembly entail the risk of over-purging, affecting genome completeness (Roach *et al.* 2018). Therefore, considering the accomplished low haplotypic duplication of the assembly and the risk of purging indispensable contigs, we removed only 141 contigs with a total sequence length of 4,186,745 bp, which were suspected artifactual by the pipeline for extraordinarily low coverage.

**Figure 2:**
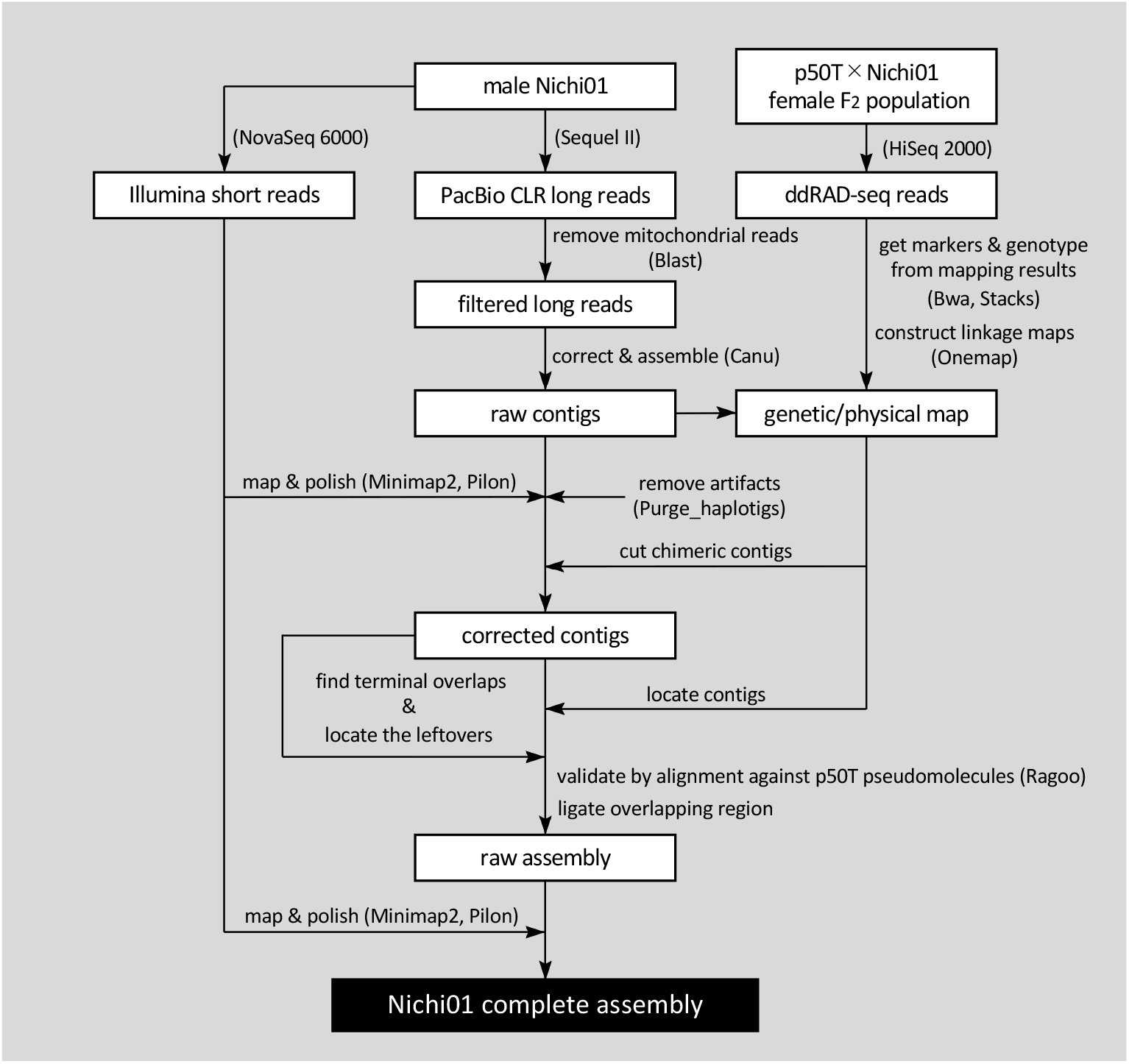
Schematic illustration of the *de novo* assembly of the Nichi01 genome. The names of the software and sequencing platforms that were used are mentioned within parentheses.

Next, to detect misassembled points in the contigs and locate them in pseudomolecules, we obtained genetic markers mapped on the contigs using ddRAD-seq reads from the F_2_ population of p50 and Nichi01. In total, 1100 markers satisfying an LOD > 3 were used for the assembly. The original classical genetic map constructed using autosomal markers contained 27 major linkage groups, and the map constructed using Z-chromosomal markers contained one major linkage group. Some genetic markers on the same contig were not included in one genetic linkage group, suggesting that the assembly contained misassembled points. Two contigs in the assembly were cut at the misassembled points detected as inconsistencies between the physical and genetic positions of the markers on the contigs and the lack of mapped read coverage at the points. Ordering and orienting the corrected contigs with the markers according to the genetic position revealed that most pairs of terminal regions of neighboring contigs overlapped over 3000 bp, indicating that the allowed mismatches in the overlapped region of the reads to be joined were excessively small. Although a more lenient assembly (less strict in joining overlapped contigs) can join more overlapped contigs, it would also induce more wrong assemblies. Therefore, we manually located the contigs missing positional markers into pseudomolecules using the overlap in the terminal region of the contigs as a clue, as long as their positions were consistent with the alignment against the pseudomolecules of p50T. Finally, we located 82 of the 380 contigs, ligated overlapping terminal regions of the neighboring contigs, and then polished the sequences again with Illumina short reads. The assembly comprised 28 pseudomolecules and 157 unplaced contigs (Table 1). The N50 value of the final assembly sequence was 16.614 Mb. All pseudomolecules except chr15, chr17, chr19, and chr24 lacked gaps in the sequence; whereas, all pseudomolecules except chr02, chr11, chr12, chr16, chr23, and chr27 contained CCTAA telomeric repeat sequences at both terminal ends (Figure 3A). Thus, 18 of the 28 chromosomal sequences were successfully assembled as telomere-to-telomere (T2T) gap-free pseudomolecules. These values have been greatly improved compared to those reported in a previous study reporting the genome assembly of p50T; only two pseudomolecules had a pair of telomeres, and none was assembled to form T2T gap-free one except for chromosome 3 (Table 1) (Kawamoto *et al.* 2019). In many cases, scaffolding by additional information for longer distance linkage is finished in a simple connection of contigs by unknown sequences represented by N, without considering the potential overlaps of contigs. This handling compromises the continuity of an assembly and leads to artificial duplications. Our results suggest that some bioinformatic tools to solve this problem, by further scaffolding using remaining internal overlaps between contigs (Hiltunen *et al.* 2021; Ruiz *et al.* 2021), or by trimming such overlapping regions from an assembly (Roach *et al.* 2018), can be helpful.

**Figure 3:**
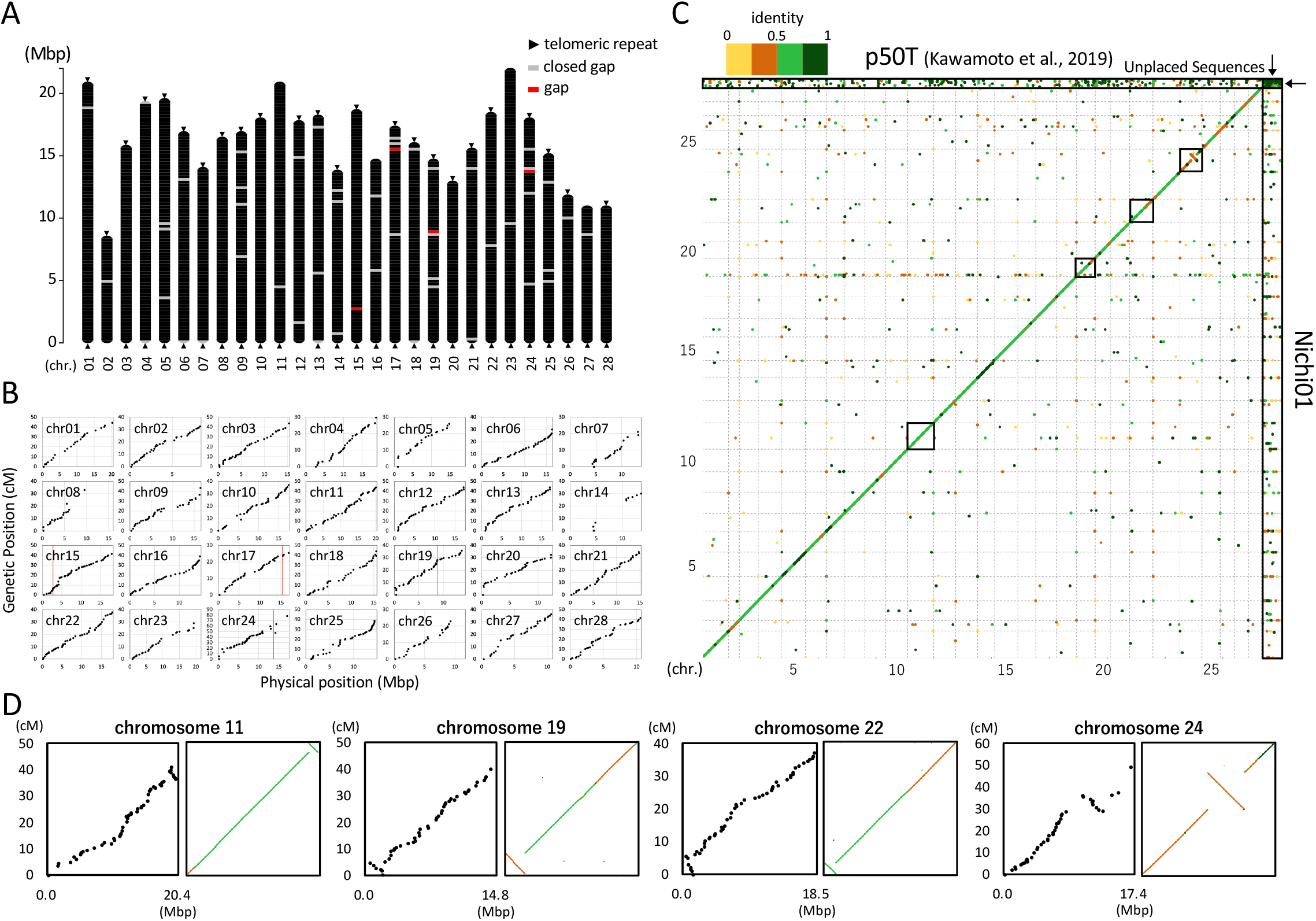
Assembly and validation of the genome of Nichi01 and p50T. (A) Illustration of pseudomolecules of the Nichi01 genome. (B) Physical and genetic position of markers on the Nichi01 pseudomolecules. The red line represents the position of the present gaps. (C) Dot-plot analysis of genome assemblies of p50T and Nichi01. Black squares highlight the pseudomolecules including structural inversion between p50T and Nichi01. (D) Physical (horizontal axis) and genetic (vertical axis) position of markers on the p50T pseudomolecules including structural inversion (left). Magnified view of the regions surrounded by black squares in (C) (right).

**Table 1:**
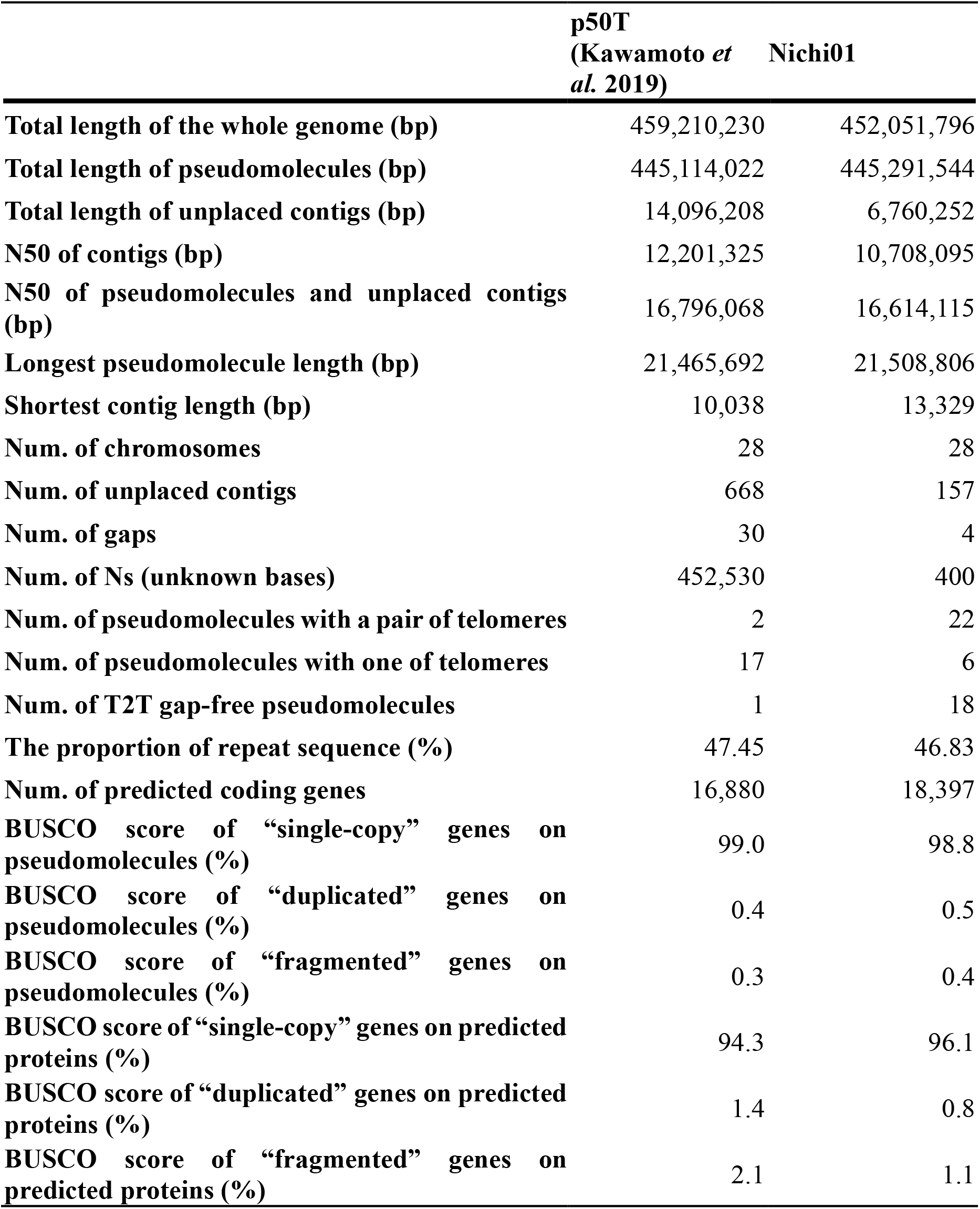
Summary statistics of the genome assembly and predicted genes of the silkworm strains, p50T and Nichi01.

|  | **p50T**<br>**(Kawamoto *et al.* 2019)** | **Nichi01** |
| --- | --- | --- |
| **Total length of the whole genome (bp)** | 459,210,230 | 452,051,796 |
| **Total length of pseudomolecules (bp)** | 445,114,022 | 445,291,544 |
| **Total length of unplaced contigs (bp)** | 14,096,208 | 6,760,252 |
| **N50 of contigs (bp)** | 12,201,325 | 10,708,095 |
| **N50 of pseudomolecules and unplaced contigs (bp)** | 16,796,068 | 16,614,115 |
| **Longest pseudomolecule length (bp)** | 21,465,692 | 21,508,806 |
| **Shortest contig length (bp)** | 10,038 | 13,329 |
| **Num. of chromosomes** | 28 | 28 |
| **Num. of unplaced contigs** | 668 | 157 |
| **Num. of gaps** | 30 | 4 |
| **Num. of Ns (unknown bases)** | 452,530 | 400 |
| **Num. of pseudomolecules with a pair of telomeres** | 2 | 22 |
| **Num. of pseudomolecules with one of telomeres** | 17 | 6 |
| **Num. of T2T gap-free pseudomolecules** | 1 | 18 |
| **The proportion of repeat sequence (%)** | 47.45 | 46.83 |
| **Num. of predicted coding genes** | 16,880 | 18,397 |
| **BUSCO score of “single-copy” genes on pseudomolecules (%)** | 99.0 | 98.8 |
| **BUSCO score of “duplicated” genes on pseudomolecules (%)** | 0.4 | 0.5 |
| **BUSCO score of “fragmented” genes on pseudomolecules (%)** | 0.3 | 0.4 |
| **BUSCO score of “single-copy” genes on predicted proteins (%)** | 94.3 | 96.1 |
| **BUSCO score of “duplicated” genes on predicted proteins (%)** | 1.4 | 0.8 |
| **BUSCO score of “fragmented” genes on predicted proteins (%)** | 2.1 | 1.1 |

### Validation of the Nichi01 genome assembly

The accuracy of the structure of pseudomolecules was validated by a consensus among the physical and genetic positions of markers, and by alignment with those of p50T (Figure 3B-D). We re-constructed the genetic map using the Nichi01 complete genome sequence and plotted the physical and genetic positions of the markers (Figure 3B); naturally, no inconsistencies were observed. However, in the results of the dot-plot analysis using p50T genome sequences, we found large inversion regions in chr11, chr19, chr22, and chr24 (Figure 3C). To verify whether they represent true structural variants or were due to a failure of either genome assembly, we examined the p50T genome assembly in detail. In the regions where the alignment-based dot-plot analysis suggested genomic sequence inversions between the two strains, an order of genetic position of the markers, which were obtained by newly mapping the ddRAD-seq reads to the p50T genome sequence, was observed to invert against an order of their physical position (Figure 3C and 3D). Additionally, between the inverted region and the other, we found gaps in the sequences of Bomo_Chr11:18,963,110-18,963,209 bp; Bomo_Chr19:21,875,89-2,187,688 bp; Bomo_Chr22:1,593, 693-1,593,792 bp; Bomo_Chr24:8,637,266-8,637,365 bp; and Bomo_Chr24:13,496,923-13,497,022 bp. Furthermore, the sequences in Bomo_Chr11:18,963,209-18,970,094 bp and Bomo_Chr22:1,593, 174-1,593,693 bp were telomeric repeats, indicating an inversion of the contigs. Thus, we concluded that sequence inversion was due to misassembly of the p50T genome sequences. We also failed to completely assemble the pseudomolecule of chromosome 24, which had been misassembled in the p50T genome. The abundant repeats of histone-like genes around 13.5 Mbp of the chromosome probably cause difficulties in assembly, which remains a future challenge.

Although the p50T genome assembly was constructed with incredible continuity, it was carried out only with 37.3 Gb of the obsolete PacBio RS II long reads and 80 BAC sequences without additional linkage information (Kawamoto *et al.* 2019). Another research group re-constructed the genome assembly of p50T with the same sequencing data and additional Hi-C, and found a discrepancy in chromosome 24 (Lu *et al.* 2019). However, it has not been validated and does not uncover the other three misassembled points. Moreover, although another *de novo* genome assembly of a different silkworm strain has been constructed previously using a combination of PacBio long-read sequencing and Hi-C, it contained many structural differences against the p50T assemblies, indicating that it contained many misscaffoldings (Tang *et al.* 2021). It is true that the Hi-C sequencing-mediated scaffolding is a convenient method and has accomplished chromosome-scale genome assemblies in many species. However, there are limitations; Hi-C data cannot accurately fix the orientation of short contigs and cause small inversions (Bickhart *et al.* 2017; Chaisson *et al.* 2015). The present improvement was accomplished by the abundant sequencing data generated by PacBio Sequel II long reads and accurate linkage information provided by over 1000 reliable positional markers. Our results suggest that a traditional genetic map is a reliable tool for providing chromosome-scale linkage information. This new reference genome with an accurate structure provides a solid research foundation for further genomics and breeding. For example, an alignment-based scaffolding using a reference genome is one of the standard methods for correction and anchoring contigs to chromosome-scale assembly; the recently published genome assembly of over 500 silkworm strains was based on the method using the p50T genome assembly (Tong *et al.* 2022). In such cases, the new Nichi01 genome assembly can serve as a better structural model.

Next, the completeness of genes in the Nichi01 genome assembly was evaluated using BUSCO (Manni *et al.* 2021) (Table 1), which identified 98.8% and 0.5% of the genes as single-copy and duplicated genes, respectively, using the reference gene set of Arthropoda. The same pipeline detected 99.0% of single-copy genes and 0.4% of duplicated genes in the p50T assembly. The BUSCO scores of both the complete or fragmented genes were 99.7% in both assemblies. All the BUSCO scores were the same when the input included unplaced contigs. These results indicate that the comprehensiveness of genes was sufficiently high and that haplotypic duplication was sufficiently low in both assemblies. Additionally, the completeness of the Nichi01 genome assembly was confirmed by the sufficiently high percentage of reads of the whole-genomic sequencing data of silkworm strains worldwide mapped to the Nichi01 genome (Supplementary Figure S4). The percentages of reads from the improved Chinese or Japanese strains were greater in Nichi01 than those in p50T, suggesting that the Nichi01 genome assembly can be a better reference genome for the NGS-based analyses of improved silkworm strains.

### Genome annotation

To annotate the assembly of repeat elements, we used RepeatMasker2 with the transposable element library provided in the Silkworm Genome Research Program (http://sgp.dna.affrc.go.jp/pubdata/genomicsequences.html) (https://www.repeatmasker.org/). As a result, a total of 211.7 Mb, representing 46.83% of the Nichi01 genome sequence, was predicted as the repeat element and included 335,620 SINEs, 258,713 LINES, 13,484 LTRs, and 47,426 DNA transposons. The frequencies of repeat elements were very similar to those observed in the p50T assembly (Kawamoto *et al.* 2019), indicating their high similarity (Figure 4). The repeat-masked sequence of the assembly was used for subsequent gene predictions. Using 30 mRNA-seq datasets from 10 different organs and curated protein sequences of silkworms registered in the RefSeq database, 18,397 genes were predicted in the Nichi01 genome (Table 1). Supplementary Table S3 shows 13,238 genes that were annotated using eggNOG-mapper (Cantalapiedra *et al.* 2021), 8,354 of which were annotated with Gene Ontology (GO) categories. The total BUSCO score of the single-copy and duplicated genes of the predicted protein sequences of Nichi01 was 96.9, which was over 1% greater than the value in p50T (Kawamoto *et al.* 2019). The increase in the number of the predicted genes and the BUSCO score in the Nichi01 genome may be due to the shorter amino acid length cutoff, the large amount of mRNA-seq used as hints, or the integration of multiple methods of gene prediction. Although this change meant improved comprehensiveness of the gene model, we note that it did not necessarily imply an improvement in the accuracy of the gene model.

**Figure 4:**
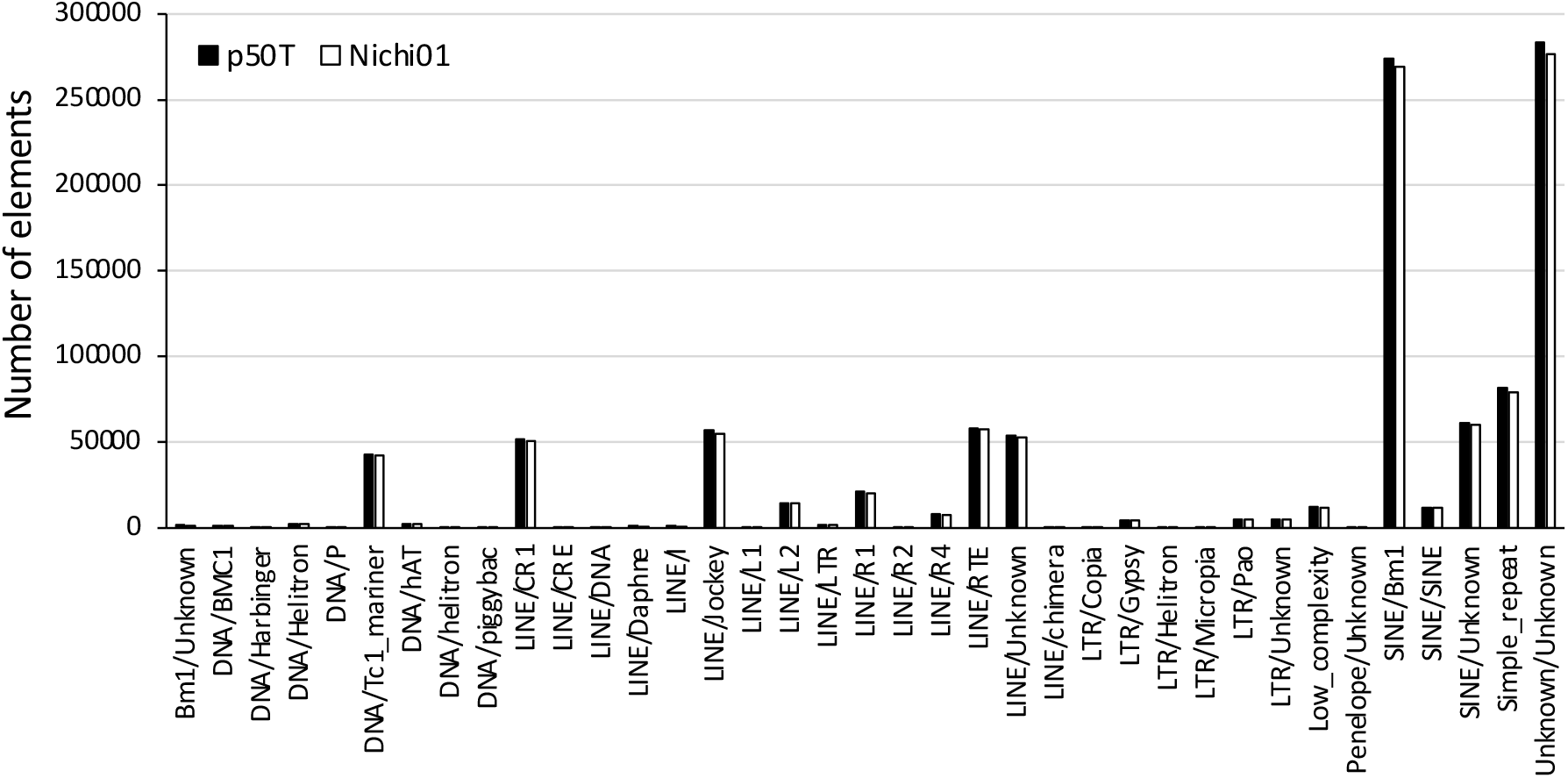
Distribution of repetitive elements in the genome of p50T and Nichi01. Repetitive elements were detected using RepeatMasker with the transposable element library available in the Silkworm Genome Research Program (http://sgp.dna.affrc.go.jp/pubdata/genomicsequences.html)

### Conclusions

This report presents the most accurate genome assembly of silkworms to date. Here, among the 28 chromosomal sequences of the improved Japanese strain Nichi01, we assembled 18 sequences as T2T gap-free pseudomolecules. In the assembly, only four gaps remained. Notably, four regions that were misassembled in the previous study on the p50T genome were corrected. Structural accuracy was confirmed using a high-density genetic linkage map. This can be considered as the gold-standard reference genome for silkworms. Moreover, we predicted genes in the assembly using over 90 Gb of mRNA-seq and published protein sequence data, resulting in an improvement in the completeness of the gene model. This study provides valuable resources for basic research on Lepidoptera and applied research on the superior protein productivity of commercial silkworms. The annotated data of the Nichi01 genome are available in KAIKObase (https://kaikobase.dna.affrc.go.jp/index.html) with a genome browser and BLAST search service.

## Data Availability Statement

The final assembly and all raw sequence data are available in the Sequence Read Archive under the BioProject accession numbers PRJDB13411, PRJDB13413, and PRJDB13956. The DDBJ accession numbers for the assembly are AP026182-AP026366. The final genome assembly and annotation files are available in KAIKObase: https://kaikobase.dna.affrc.go.jp/index.html, hosting the BLAST search service and the genome browser. Supplementary materials below are available at Figshare: https://doi.org/10.6084/m9.figshare.21443007.v1. Supplementary figure S1: Silk gland of Nichi01 fifth-day final instar larva; supplementary figure S2: Coverage histogram of the PacBio long reads on the corrected contigs; supplementary figure S3: Explanatory drawing of misassembly correction; supplementary figure S4: Proportion of mapped genomic short reads; supplementary table S1: The list of the published whole-genomic sequencing data used in the analysis; supplementary table S2: Information of the original sequencing data used in the analysis; supplementary table S3: Annotation result of the predicted genes in Nichi01 by eggNOG-mapper.

## Acknowledgments

We thank Kaoru Nakamura, Toshihiko Misawa, and Koji Hashimoto for rearing the silkworms. The computation was partly performed using the supercomputer of AFFRIT, MAFF, Japan. We thank Hiroki Sakai and Shuichiro Tomita for their valuable advice.

## Conflict of Interest

The authors declares that the research was conducted in the absence of any commercial or financial relationships that could be construed as a potential conflict of interest.

## Funder Information

This study was supported by the Ministry of Agriculture, Forestry, and Fisheries.

---

**Supplementary Figure S1:** Silk gland of Nichi01 fifth-day final instar larva.

Dotted red lines indicate cut points in sampling.

**Supplementary Figure S2**: Coverage histogram of the PacBio long reads on the corrected contigs.

A coverage histogram was generated using Purge Haplotigs (Roach *et al.* 2018). A total of 41.6 Gb of randomly sampled raw reads were used as the input.

**Supplementary Figure S3**: Explanatory drawing of misassembly correction.

**Supplementary Figure S4**: Proportion of mapped genomic short reads.

Gray: result in mapping to the p50T genome (Kawamoto *et al.* 2019), White: result in mapping to the Nichi01 genome. The labels were based on a previous report (Xiang *et al.* 2018). JPN_L: Japanese local strain (6 samples); JPN_I: Japanese improved strain (16 samples); CHN_L: Chinese local strain (30 samples); CHN_I: Chinese improved strain (15 samples); IND_L: southern Indian strain (3 samples); WILD: wild silkmoth *Bombyx mandarina* (6 samples). Information on used sequencing data was summarized in Supplementary Table S1. The *P*-values were calculated using the Wilcoxon rank sum test.

## Notes

https://doi.org/10.6084/m9.figshare.21443007.v1

### Competing Interest Statement

The authors have declared no competing interest.

